# Pregnancy History and Estradiol Influence Spatial Memory, Hippocampal Plasticity, and Inflammation in Middle-aged Rats

**DOI:** 10.1101/2024.06.21.600090

**Authors:** Tanvi A. Puri, Stephanie E. Lieblich, Muna Ibrahim, Liisa A. M. Galea

**Author notes:** = Corresponding Author: Dr. Liisa Galea, Treliving Family Chair in Women’s Mental Health Centre for Addiction and Mental Health, 250 College Street Toronto, ON, M5T 1R8, Canada.

## Abstract

Pregnancy and motherhood (parity) can have long-term effects on cognition and brain aging in both humans and rodents. Estrogens are related to cognitive function and neuroplasticity. Estrogens can improve cognition in postmenopausal women, but the evidence is mixed, in part due to differences in hormone therapy dose and composition. In addition, past pregnancy influences brain aging and cognition, with earlier age of first pregnancy being associated with poorer outcomes with aging. However, few studies have examined specific features of pregnancy history such as the age of first pregnancy or the possible mechanisms underlying these changes. We examined whether maternal age at first pregnancy and estradiol treatment differentially affected hippocampal neuroplasticity, inflammation, activation, and cognition in middle-age. Thirteen-month-old rats (who were nulliparous (never mothered) or previously primiparous (had a litter) at 3 months or 7 months) received daily injections of estradiol (or sesame oil vehicle) for sixteen days and were tested on the Morris Water Maze. An older age of first pregnancy was associated with impaired spatial memory but improved performance on reversal training, and increased new neurons in the ventral hippocampus compared to the other groups. Estradiol decreased total activation and percent activation of new neurons in the dorsal hippocampus, regardless of parity history. Estradiol also decreased the production of anti-inflammatory cytokines based on age of first pregnancy. This work suggests that estradiol affects neuroplasticity and neuroinflammation in middle age, and that pregnancy history can have long lasting effects on hippocampus structure and function.

## 1. Introduction

Pregnancy and childbirth can have long-lasting effects on the maternal brain (Duarte-Guterman et al., 2019; Puri et al., 2023). Significant changes take place in the body during pregnancy and the postpartum, including drastic modifications to the endocrine and immune systems (Brett & Baxendale, 2001; Haim et al., 2017). The brain also undergoes substantial changes in both structure and function during pregnancy, and some of these changes persist into middle age or later (de Lange et al., 2020; Duarte-Guterman et al., 2019; Orchard et al., 2020, 2021; Pawluski et al., 2022; Puri et al., 2023).

In humans, brain regions such as the frontal cortex and hippocampus show reduced volume after pregnancy that is evident at least six years later (Hoekzema et al., 2017; Martínez-García et al., 2021). Rodents experience similar changes with pregnancy, as primiparous rats (that have been pregnant and mothered once) show decreased hippocampal volume and reduced neurogenesis in the hippocampus during gestation and into the late postpartum (Eid et al., 2019; Galea et al., 2000). In middle age, however, reproductive experience is associated with less evident brain aging (as measured by grey matter volume) in humans, and increased hippocampal neurogenesis and the synaptic protein PSD-95 in rodents (Barha et al., 2015; de Lange et al., 2019, 2020; Duarte-Guterman et al., 2023; Eid et al., 2019; Galea et al., 2018; Puri et al., 2023). These findings show that the hippocampus displays dynamic plasticity from the short to the long term with reproductive experience.

Pregnancy history is associated, however, with mixed findings on cognition in middle age, and conflicting data may be due to both the type of cognition tested and pregnancy history characteristics such as amount of parity and age of first pregnancy (Colucci et al., 2006; Cui et al., 2014; Gatewood et al., 2005; Jang et al., 2018; Puri et al., 2023). Earlier ages of first pregnancy show less beneficial effects on cognition and increased dementia risk later in life compared to later ages of first pregnancy (Almanza-Sepulveda et al., 2018; Chico et al., 2014; Fu et al., 2023; Gong et al., 2022; Rocca et al., 1991). Although there is clear evidence pregnancy leaves long-term signatures on the brain and on behavior, pregnancy characteristics, such as the age of first pregnancy, remain largely understudied.

Reproductive experience alters inflammation in the brain in both the short and long-term (Eid et al., 2019; Haim et al., 2017). In rats, maternal experience increases pro-inflammatory and decreases anti-inflammatory cytokine expression, and alters microglial morphology such that they become more ameboid in the early postpartum period (Duarte-Guterman et al., 2023; Eid et al., 2019; Haim et al., 2017). In middle age, primiparity increases the concentrations of IFN-γ and IL-4 but decreases the concentration of IL-1β peripherally in middle-aged rats (Eid et al., 2019). Primiparity reduces inflammatory signaling in the hippocampus compared to nulliparity (never pregnant or mothered) in wildtype middle-aged rats (Lee et al., 2023). Therefore, motherhood has a lasting effect on inflammation in the periphery and in the brain (Bourgognon & Cavanagh, 2020; Lee et al., 2023; Villa et al., 2018). The body’s immune cells secrete cytokines to regulate various functions, including neuroplasticity (Villa et al., 2018) and cognition (Bourgognon & Cavanagh, 2020), but how these cytokines could be related to cognition with age of first pregnancy is not yet known, and will be investigated in this study.

Estradiol treatment in middle aged females affects neuroplasticity, inflammation, and cognitive function depending on cognitive task, dose, and previous reproductive experience (Barha et al., 2015; Barha & Galea, 2011; Galea et al., 2018; Luine, 2014; McCarrey & Resnick, 2015; Shifren et al., 2019). In humans, treatment with estradiol-based menopausal hormone therapies (MHTs) improves cognition and reduces risk for neurodegenerative diseases in middle age (Hogervorst et al., 2000; Kim et al., 2021; McCarrey & Resnick, 2015; Wharton et al., 2011), but these effects depend on formulation, dose, route of administration, and timing of treatment onset relative to menopause (Craig et al., 2005; Espeland et al., 2013; Kim et al., 2021; Resnick et al., 2009). How estrogens affect cognition, inflammation, and neuroplasticity factoring parity history is less well understood.

Estrogens can have differential effects on neuroplasticity depending on parity history in middle age. Acute estrogens increase hippocampal cell proliferation in multiparous middle aged rats, but not in age-matched nulliparous controls (Barha & Galea, 2011). Chronic estrogens (22 days) decrease neurogenesis regardless of parity, depending MHT formulation or sub-region of the dentate gyrus (Barha et al., 2015; Galea et al., 2018). To the best of our knowledge, there is only one study that has investigated the effects of estrogens on inflammation across previous parity. Premarin (an estrone-based MHT) increased serum TNFα in nulliparous rats, but decreased it in primiparous rats (Galea et al., 2018). Together, these data indicate that previous reproductive experience and estrogens influence neuroplasticity, cognition, and peripheral inflammatory signaling, but how age of first pregnancy influences these findings in middle age is not yet known and this study aims to fill that gap.

The aim of our study was to investigate how parity history (age of first pregnancy) affects brain function, inflammation, and neurogenesis in middle aged rats. We hypothesized that both the age of first pregnancy and treatment with estradiol would affect spatial memory performance, and parity would increase neurogenesis in middle aged rats. We also hypothesized that estradiol would alter the activation of new neurons in the hippocampus and cytokine concentration dependent on previous reproductive experience.

## 2. Methods

### 2.1 Animals

Eighty one female and twenty male Sprague-Dawley rats were purchased from Charles River (Quebec, Canada) weighing 200-250g (2 months old) upon arrival at the University of British Columbia. Rats were maintained on a 12:12 light:dark cycle (lights on at 0700h) in translucent polyurethane bins (24 x 16 x 46 cm) under standard laboratory conditions (21±1°C, 50±10% humidity) and had *ad libitum* access to standard laboratory chow (Jamieson’s Pet Food Distributors, Canada) and tap water. All rats were pair housed except during the breeding period. Nulliparous females were pair housed except when housed singly, and for an equivalent time to primiparous females, in a separate colony room from the primiparous rats during breeding and pup rearing to avoid exposure (odors and sound) from pups and males. Apart from breeding procedures, rats were left undisturbed except for weekly cage changes until they reached testing age at 13 months old. All procedures were performed in accordance with the ethical guidelines set forth by the Canadian Council on Animal Care and approved by the Animal Care Committee at the University of British Columbia. All efforts were made to reduce the suffering and number of animals.

### 2.2 Breeding procedure

Rats were randomly assigned to one of three groups: nulliparous (never pregnant and with no pup experience, n = 24), younger dams (rats that were pregnant at 3-4 months old, n= 25), or older dams (rats that were pregnant at 7-8 months old, n=31) (see Figure 1 for experimental timelines). At 1800h daily, 2 females and 1 male were pair housed together overnight. If sperm cells were detected the next morning in lavage samples, females were considered pregnant and at Gestation Day 0 (GD0). GD0 rats were single housed in a clean Allentown cage with aspen chip bedding and left undisturbed until parturition except for weekly cage changing and weighing. If sperm was not detected, females were paired with a male rat daily until sperm cells were detected or 20 days had passed. In total, eighteen of the fifty six rats did not get pregnant most of which were from the older age of first pregnancy group; sixteen (92%) were from the older age group and two were from the younger age group (8%). These rats were excluded from all further experiments.

**Figure 1.**
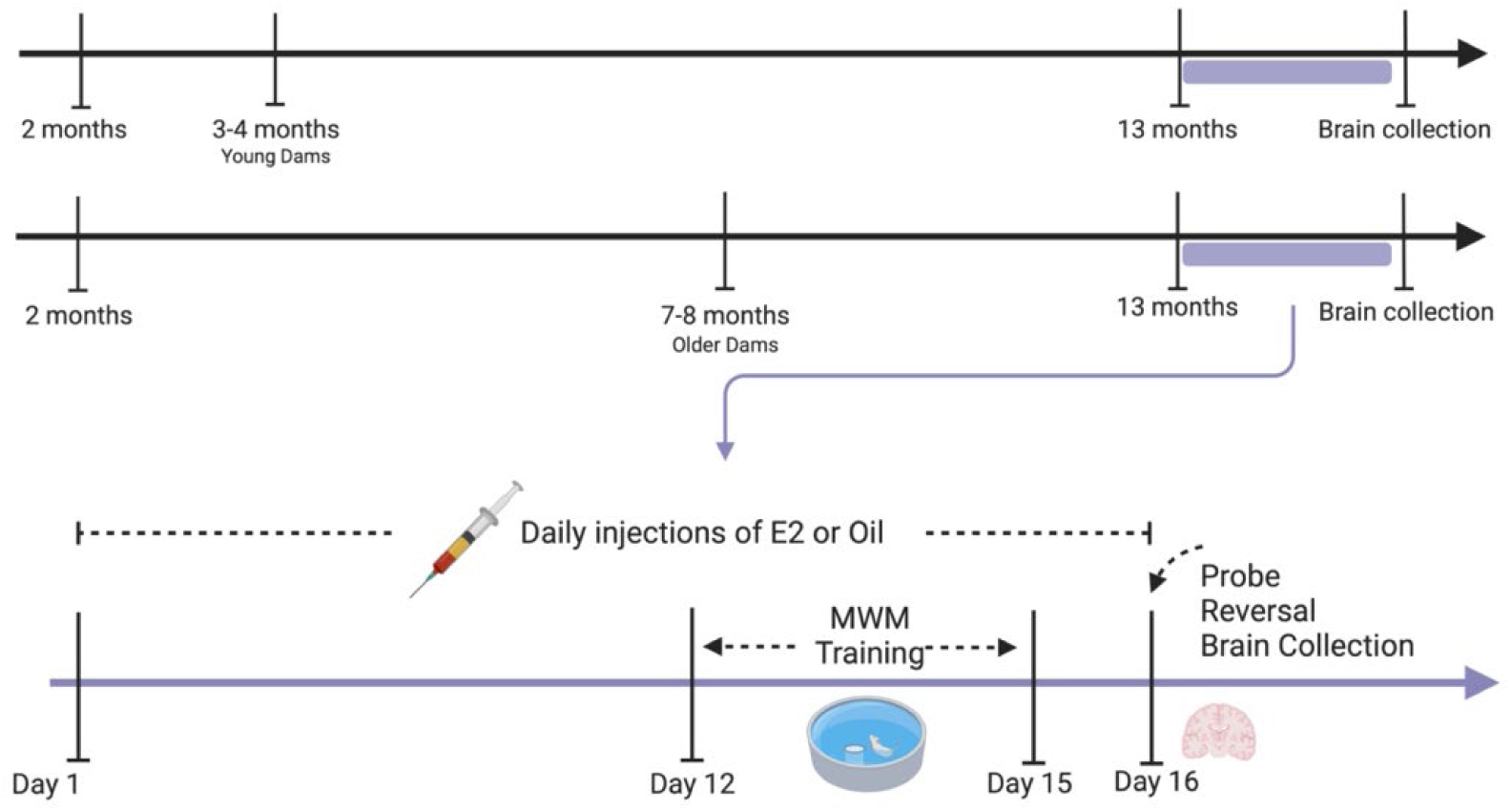
Experimental timeline: Female rats were either bred at 3-4 months old, at 7-8 months old, or were nulliparous. At middle age (13 months) rats received a daily injection of E2 or oil vehicle for 16 days. On days 12-15, rats were trained on a standard reference memory version of the MWM. A probe trial and reversal trials were performed on Day 16, followed by brain collection. MWM = Morris water maze, E2 = estradiol. Figure made using BioRender.com.

On Postpartum Day (PPD) 1 all litters were culled to 4 males and 4 females. If the number of pups or sex ratio was insufficient in a litter, pups were cross fostered from a dam that had given birth the same day, when possible. Pups were weaned at PPD 23, and dams returned to a pair-housed condition at PPD 30.

### 2.3 Maternal behavior observations

Maternal behaviors were recorded twice a day from PPD2 to PPD8. Dams were recorded for 30 minutes using small, unobtrusive cameras (PapaKoyal Spy Camera, Amazon.ca) from 8:00-8:30am and 5:00-5:30pm. The amount of time that dams spent engaging in licking, grooming, nursing (passive, blanket, and arch-back), retrieving pups, nest building, and off-nest behaviors (eating, drinking, self-grooming, and sleeping off nest) for 10 continuous minutes was then quantified from the videos. The ratio of how much time the dams spent conducting on-nest: off-nest behaviors was also calculated.

### 2.4 Hormone administration

At 13 months old, animals received a subcutaneous injection of either estradiol (0.3ug in 0.1mL sesame oil, 17β-estradiol, Sigma, St. Louis, MO, USA) or an equivalent volume of sesame oil vehicle (Sigma, St. Louis, MO, USA) every day until experimental endpoint (16 days). The dose of estradiol was chosen based on previous studies that it enhanced cell proliferation in young adult female rats, and adjusting for weight, it is also a dose frequently prescribed to menopausal individuals (Barha et al., 2009; Corbelli et al., 2015; Studd et al., 1999). Injections were administered between 8:00 and 9:30am and at least 2 hours before the start of any behavioral testing.

### 2.5 Estrous Cycle

Estrous cycle stages were monitored daily for the duration of hormone injections and behavior, as the proestrus stage alters both cell proliferation and spatial learning (Korol et al., 2004; Tanapat et al., 1999). Rats were lavaged every day following hormone injections by inserting a small glass dropper with 200-500μL of water into the vagina and collecting a sample. The sample was then transferred to a slide and stained with Cresyl Violet (Sigma, Oakville, Canada) and examined under a microscope at 20x in white light. Estrous state (proestrus, estrus, metestrus, or diestrus) was determined using previously published guidelines (Ajayi & Akhigbe, 2020; Cora et al., 2015). If animals consistently cycled through at least three of four stages in 5 days, they were classified as ‘cycling’, otherwise as ‘irregular’.

### 2.6 Morris Water maze (MWM)

#### 2.6.1 Apparatus

The Morris water maze used was a circular pool 180cm in diameter, filled 60cm deep. The inside of the pool was painted black and filled with room temperature water mixed with non-toxic tempera paint to render it opaque. Large and distinct cues were placed on each of the four walls of the room and red light lamps were placed in two corners. Both were unchanged throughout behavioral training. A square platform 10cm wide was submerged approximately 2cm below the surface of the water such that it was invisible from above. Measures used to track the animals in the pool were collected using cameras installed on the ceiling connected to ANY-maze 7.10 software (Stoelting, Wood Dale, IL, USA). ANY-maze was used to capture several variables of interest, including latency and distance to reach the platform, as well as the percentage of time spent in each quadrant and in the platform zone.

#### 2.6.2 Training

*Visible platform training:* Initially, rats were put in the Morris water maze to find a visible platform elevated 2cm above the water level. If the rat did not find the platform within 60s, the experimenter guided it to the platform and it was allowed to remain there for 15s before being retrieved and returned to the home cage. If a rat did not find the visible platform in three consecutive trials, it was excluded from further behavioral testing. Four rats could not find the platform at all during this phase of testing (two nulliparous, one younger dam, and one older dam, all oil treated). These animals were classified as visually impaired and were excluded from any further testing.

*Hidden platform training:* Rats received 4 days of training on the standard reference-memory version of the MWM. Subjects underwent 4 trials per day, where they had 60 seconds to find the submerged platform (that remained in the same place for all training days and trials). If the rat did not find the platform, the experimenter guided it to the platform and allowed it to remain there for 15s before returning it to the home cage. Inter-trial interval was 5 minutes. Within each session, the rat was put in the maze from a pseudo-random cardinal compass point each day, and the same starting point was not repeated for any given rat on one day. Any debris floating in the water was removed between every trial, and the entire pool drained, cleaned, and refilled every other day.

#### 2.6.3 Probe Trial and Reversal Learning

On the 5^th^ day of MWM trials, rats received a 60s probe trial, in which there was no platform in the maze, to measure memory retention of the original platform location. To determine how well rats recalled the location of the previously hidden platform, we analyzed the percentage of time rats spent in the target quadrant.

*Reversal learning:* Twenty minutes after the probe trial, the hidden platform was reintroduced into the MWM, but in the opposite quadrant than the previous trials. Rats performed 4 trials, with an inter-trial interval of 5 minutes. This trial was used to investigate how efficiently rats would learn a new location for the hidden platform.

### 2.7 Tissue collection and processing

Animals were euthanized 90 minutes after the probe trial of the MWM by decapitation. Brains were collected and bisected along the sagittal plane. The left hemisphere of the brain was flash frozen on dry ice, then frozen at –80°C until it was used for cytokine evaluation using electro-chemiluminescent assays. The right hemisphere of the brain was postfixed in freshly prepared cold 4% paraformaldehyde solution for 24 hours, and then placed in a 30% sucrose in 0.1M phosphate buffer saline solution (PBS, pH = 7.4) at 4°C until brains sank.

The flash-frozen left half of the brain was coronally sliced at 300μm using a cryostat (CM3050 S, Leica, Wetzlar, Germany) at –15°C. The dorsal and ventral hippocampus were identified and dissected using tissue punching tools (0.35mm, 0.75mm, and 1.20mm diameter, Harris Uni-Core, Sigma-Aldrich) on dry ice, and placed into tubes containing 10 lysing beads (1.4mm ceramic spheres, Lysing Matrix D, MP Biomedicals, Santa Ana, CA, USA). The tissue was then homogenized in 120μL Complete lysis buffer (Meso Scale Discover, Rockville, MD, USA) using the Qiagen TissueLyserII (Qiagen, Hilden, Germany). After homogenization, samples were centrifuged at 1200g for 10 minutes at 4°C, and supernatants were stored at –80° until cytokine analysis.

The post-fixed right half of the brain was sliced for immunohistochemistry. Brains were rapidly frozen using dry ice, and coronal sections of 30μm were collected from approximately bregma 3.72mm and –6.96mm (Paxinos & Watson, 2014) using a freezing microtome (Leica, Richmond Hill, ON, Canada). The dorsal and ventral hippocampus (bregma –2.64mm and – 6.36mm) were collected in 10 series (Paxinos & Watson, 2014). Sections were stored in anti-freeze solution (30% ethylene glycol, 20% glycerin in 0.1M PBS, pH = 7.4) at –20°C until used for immunohistochemistry.

### 2.8 Brain cytokine quantification

Cytokine levels in the dorsal and ventral hippocampus were quantified using a multiplex electrochemiluminescence immunoassay kit (V-PLEX Proinflammatory Panel 2, Rat) from Meso Scale Discovery (Rockville, MD, USA). The panel quantified the following cytokines and chemokines in each sample: interferon gamma (IFN-γ), interleukin (IL)-1β, IL-4, IL-5, IL-6, IL-10, IL-13, tumor necrosis factor (TNF)-α, and C-X-C motif ligand 1 (CXCL1). All samples were run in duplicate, and intra-sample variability was <20%. All plates were read using the Sector Imager 2400 (Meso Scale Discovery), and data acquired and analyzed using the Methodical Mind and Discovery Workbench 4.0 software (Meso Scale Discovery). This assay contains cytokines that are considered proinflammatory (IL-1β, IFN-γ, TNF-α), pleiotropic (IL-6), and anti-inflammatory (IL-4, IL-10, IL-13). These individual markers can provide a broad view of the inflammatory state of the brain, and can provide a comprehensive evaluation of the cytokine milieu in brain tissue.

Total protein levels were quantified in tissue homogenates in triplicates using the Pierce Microplate BCA Protein Assay Kit (Thermo Scientific Pierce Protein Biology, Thermo Fisher Scientific, Waltham, MA, USA). Individual cytokine levels are reported as pg/mg of protein in each sample.

### 2.9 Doublecortin and cFOS Immunohistochemistry

Doublecortin is a microtubule binding protein that is expressed in immature or newly born neurons for 21 days in adult rats, and therefore was used as a marker for immature neurons in the dentate gyrus (Brown et al., 2003). c-FOS is the protein of the immediate-early gene *c-fos* which is transiently expressed in neurons in response to action potential and used as a marker of cellular activation (reviewed in (Kovács, 2008)).

Briefly, coronal sections were washed 4x in 0.1M PBS for 10 minutes per wash at room temperature, then washed with 10% Triton-X in 0.1M PBS (Sigma Aldrich, St. Louis, MO, USA) for 30 minutes. Sections were then blocked in 10% Normal Donkey Serum (NDS), 0.3% Triton-X in 0.1M NDS, followed by a double overnight incubation at 4°C in Doublecortin and cFOS primary antibodies (1:1000 anti-Doublecortin goat polyclonal IgG, sc-8066, Santa Cruz Biotechnology, Dallas, TX, USA; 1:1000 anti-cFOS rabbit Ab, 190289, Abcam, Toronto, ON, Canada) in 10% NDS and 0.3% Triton-X in 0.1M PBS. Two days later, sections were washed 3x in 0.1M PBS for 10 minutes per wash at room temperature, then incubated with secondary antibodies (1:500 Alexa488 donkey anti-goat, A11055, ThermoFisher, Waltham, MA, USA; 1:500 Alexa594 donkey anti-rabbit, A21207, ThermoFisher) in 10% NDS and 0.3% Triton-X in 0.1M PBS for four hours. Sections were washed 3x in 0.1M PBS for 10 minutes and then counterstained with DAPI for 2.5 minutes (14.3mM solution diluted 1:5000 in 0.1M PBS, ThermoFisher). Finally, cells were washed 3×5 minutes and then mounted on Superfrost Plus slides (Fisher Scientific, Inc., Hampton, NH, USA), and cover-slipped with anti-fade medium (0.5% polyvinyl alcohol-DABCO, Sigma, St. Louis, MO, USA).

### 2.10 Microscopy

All quantification of protein expression was conducted by an experimenter blind to the group assignment of each animal. Slides were scanned using the Zeiss Axio ScanZ1 (Carl Zeiss Microscopy, Thornwood, NY, USA) at 20x using an air immersion objective lens, and manually counted using Zen 3.0 software to visualize sections (Carl Zeiss Microscopy, Thornwood, NY, USA) on digitized images from every 10^th^ section. Sections were categorized as dorsal or ventral dentate gyrus using established criteria (Banasr et al., 2006). Cells were counted separately in each region, as they each have specialized function: the dorsal hippocampus is associated more with spatial learning and reference memory and the ventral hippocampus is more associated with stress and mood (Kjelstrup et al., 2002; Moser et al., 1993).

All counts were completed on two dorsal and two ventral sections of the dentate gyrus. Doublecortin-immunoreactive (DCX-ir) cells in the granule cell layer were counted. The percentage of activated new neurons was calculated by exhaustively identifying how many of the DCX-ir cells were co-expressing cFOS. The total number of activated cells was calculated by exhaustively manually counting cFOS-ir cells in the dentate. Of the percent activated neurons, two outliers (>2 standard deviations) were removed (both estradiol treated, one pregnant at a younger age and the other nulliparous in the dorsal hippocampus). Areas of the dorsal and ventral dentate gyrus used for protein quantification were calculated using ImageJ.

### 2.11 Data Analyses

Data were analyzed using multifactorial analysis of variance (ANOVA), with maternal age (younger dams, older dams, and nulliparous rats) and hormone therapy (E2 or vehicle) as between-subject factors. Performance on the training of the MWM was analyzed using a repeated measures ANOVA with training day (1-4) as within-subject variables. The number of DCX-ir, cFOS-ir, and percent of double-labeled cells were analyzed using a repeated measures ANOVA with region (dorsal and ventral hippocampus) as a within-subjects factor, and between-subjects factors as described above. Cytokines were analyzed individually with region (dorsal and ventral hippocampus) used as the within-subjects factor. Effect sizes (partial η^2^ or Cohen’s *d*) are reported for significant effects as appropriate. Post-hoc comparisons used Newman-Keuls and any *a priori c*omparisons were subjected to Bonferroni correction. All data for ANOVAs were analyzed using Tibco Statistica (v.9, StatSoft, Inc., Tulsa, OK, USA). Chi-square analyses were performed with R and RStudio, using base R functions (script may be obtained by contacting the corresponding author; R version 4.3.1). Significance level of *p* ≤0.05 was used, and trends discussed if 0.05 ≤ p ≤ 0.10.

## 3. Results

### 3.1 Older Dams have fewer pups than younger dams. Estradiol decreased cycling regularity regardless of reproductive experience

As mentioned earlier, a much lower proportion of older rats successfully carried pregnancies to term compared to younger rats, as 48% of older rats (15/31) bred successfully, whereas 92% of younger rats (23/25) bred successfully [χ^2^ (1) = 12.069, p < 0.001]. Rats pregnant at an older age had fewer pups than rats pregnant at a younger age [Main effect of age of first pregnancy: F (1,48) = 39.484, p < 0.001, partial η^2^ = 0.45; Appendix Figure A.1A]. However, there was no significant effect of age of first pregnancy on maternal behavior (all p’s > 0.11; see Appendix Table A.1).

Rats treated with estradiol were less likely to be cycling compared to animals treated with oil vehicle [χ^2^ (1) = 6.760, p = 0.009; Figure A.1b]. There was no significant effect of parity or age of first parity on the likelihood of having a regular estrous cycle at 14 months old (all p’s > 0.280). There was a trend towards an effect of pregnancy history on body weight [main effect of pregnancy: F (2,83) = 2.642, p = 0.077, partial η^2^ = 0.060], with nulliparous rats weighing more than previously parous rats (Appendix Figure S1c). There were no group differences in adrenal weight (all p’s > 0.23).

### 3.2 An older age of first pregnancy improved spatial memory, but did not significantly influence acquisition in the Morris water maze

There were no significant main or interaction effects on average time to reach the visible platform (all p’s >0.095). However, four rats could not find the visible platform, and these animals were excluded from any further testing (all oil treated; two nulliparous, one younger dam, one older dam).

As expected, rats swam a shorter distance and took shorter latencies to reach the hidden platform across days during the Morris Water maze training trials [distance: main effect of day: F (3,153) = 28.194, p < 0.001, partial η^2^ = 0.356; Figure 2A and 2B; latency: main effect of day: F (3,159) = 36.734, p < 0.001, partial η^2^ = 0.409, Figure 2D and 2E]. There were no other significant main or interaction effects on distance or length of time rats swam to the hidden platform during the acquisition phase (all p’s > 0.25).

**Figure 2.**
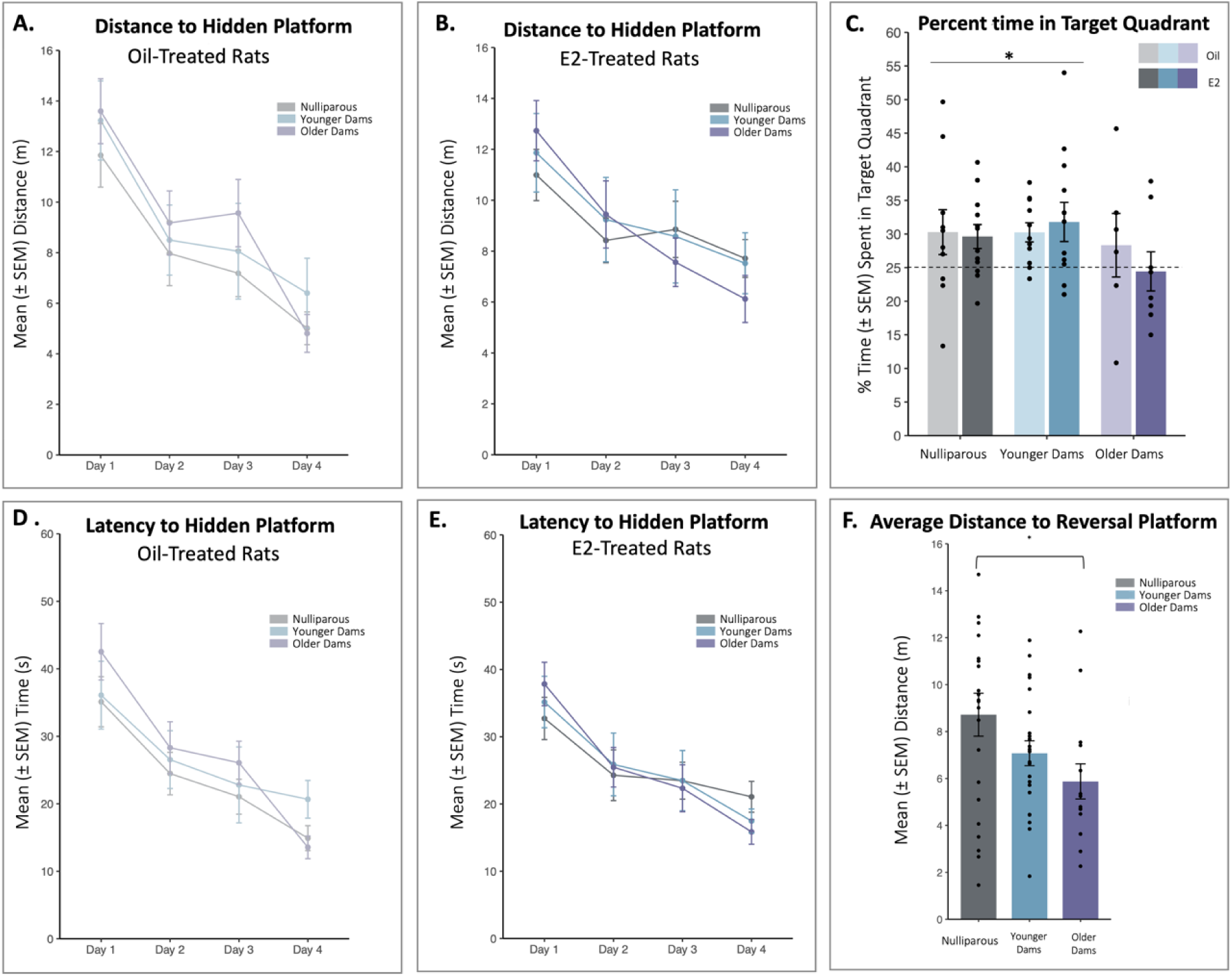
Distance (A, B, F) and latency (D, E) data from the acquisition and (C) probe trials of the MWM. (A-B) Group means ± SEM (standard error of the mean) of the distance (A) oil-treated and (B) E2-treated animals swam to find the hidden platform. (D-E) Group means ± SEM of the time (D) oil-treated and (E) E2-treated animals swam to find the hidden platform. Rats found the hidden platform faster each day. (C) Percent of time rats spent in the target quadrant during the probe trial of the MWM. (F) Mean ± SEM average distance animals swam to the hidden platform for all four reversal trials grouped by age of first pregnancy. MWM = Morris water maze; E2 = estradiol.

To examine whether rats recalled the position of the hidden platform during the probe trial, we examined the percent of time animals spent in the target quadrant compared to chance. All groups performed above chance except the older dams, regardless of estradiol treatment (Older dams p’s >0.5; nulliparous: oil p = 0.052 one tailed, estradiol p = 0.025, younger dams: oil p = 0.004, estradiol p = 0.040, Figure 2C).

### 3.3 Dams pregnant at an older age showed enhanced reversal training

All rats travelled shorter distances to reach the new platform location across trials, with significantly shorter distances on Trial 3 or Trial 4 compared to Trial 1 (p < 0.001, Cohen’s *d* = 0.930) and 2 (p = 0.001, Cohen’s *d* = 0.631; Trial 3 compared to Trial 1 (p < 0.001, Cohen’s *d* = 0.634) and 2 (p = 0.043, Cohen’s *d* = 0.346) [main effect of trial: F(3,159) = 8.789, p < 0.001, partial η^2^ = 26.368; Figure 2F]. A priori we expected differences with age of first pregnancy, and we found that dams that were pregnant at an older age swam shorter distances than nulliparous rats (p = 0.017, Cohen’s *d* = 0.500) but not younger dams (p=0.11) [main effect of pregnancy: F (2,53) = 2.881, p = 0.065, partial η^2^ = 0.098].

### 3.4 Older dams had more immature neurons in the ventral hippocampus than younger dams

As there were no group differences in the areas from which cells were counted, estimated counts are used (all p’s > 0.20). As expected, there were more DCX-ir cells in the ventral compared to the dorsal dentate gyrus regardless of the age of first pregnancy or estradiol treatment [main effect of region: F (1,39) = 89.012, p = <0.001, partial η^2^ = 0.695; Figure 3A and 3B]. There was a trend towards an interaction between HPC region and reproductive history, and a priori analyses indicated that older dams had more ventral DCX-ir cells than younger dams (p=0.006, Cohen’s *d* = 0.756) but were not statistically different from nulliparous rats (p = 0.054, did not survive Bonferroni Correction). [Region by pregnancy status interaction: F (2,39): 2.716, p = 0.079, partial η^2^ = 0.0.122]. There were no other main effects or interactions (all p’s > 0.28).

**Figure 3.**
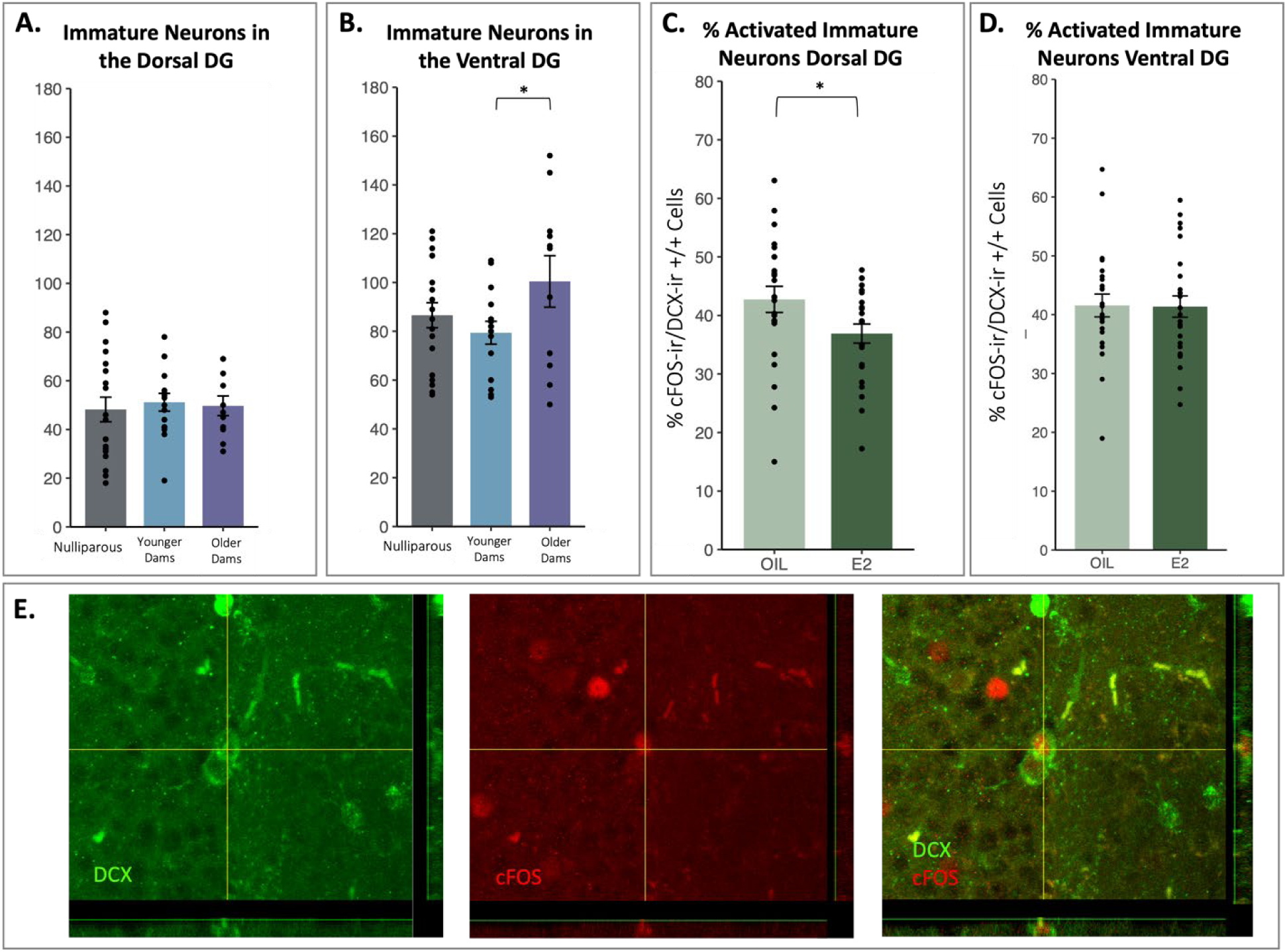
(A-B) Mean ± SEM (standard error of the mean) of the number of DCX-ir cells in the dorsal and ventral dentate gyrus. Ventral DG had more DCX-ir cells than dorsal DG regardless of age of first pregnancy or treatment. (B) A priori: Dams pregnant at an older age had more immature neurons in the ventral hippocampus than younger dams. (C-D) Mean ± SEM (standard error of the mean) of the percent of DCX-ir cells in the dentate gyrus that were also expressing cFOS. (C) Estradiol decreased the percent activation of immature neurons in the dorsal DG graphed by treatment. (D) There was no effect of estradiol or parity on the percent activation in the ventral dentate gyrus. (E) Photomicrographs of cells stained with DCX, cFOS, and co-labeled with both. * Indicates p < 0.05. 10 Abbreviations: DG = dentate gyrus; E2 = estradiol.

### 3.5 Estradiol decreased the percent of activated immature neurons in the dorsal dentate gyrus

To determine whether estradiol and age of first pregnancy affected activation of new neurons in the dentate gyrus after spatial memory retrieval we analyzed the percentage of DCX-ir cells that co-expressed cFOS. Estradiol decreased the percentage of double-expressing cFOS-ir and DCR-ir cells in the dorsal (p=0.004, Cohen’s *d* = 0.640), but not ventral dentate gyrus (p=0.866), regardless of parity [treatment by region interaction: F (1,41) = 4.901, p = 0.032, partial η^2^ = 0.106; Figure 3B]. There were no other significant main effects or interactions (all p’s > 0.16).

### 3.6 Older dams had more activation in the ventral DG, which was lowered with estradiol

In the ventral DG, older dams had greater overall cFOS expression than all other groups (all p < 0.04). Additionally, estradiol decreased activation compared to oil in older dams (p < 0.001, Cohen’s *d* = 1.765), but estradiol had no significant effect in the other groups (younger dams (p = 0.88), nulliparous (p = 0.96) [interaction of region by treatment by age of first pregnancy: F (2,42) = 4.905, p = 0.012, partial η^2^ = 0.189; Figure 4B]. There was also a main effect of region [F (1,42) = 31.309, p < 0.001, partial η^2^ = 0.629; Figure 4A and 4B], and a main effect of treatment [F (1,42) = 3.9226, p = 0.05, partial η^2^ = 0.0854; Figure 4A]. There were no other significant effects of interactions (all p’s > 0.160).

**Figure 4.**
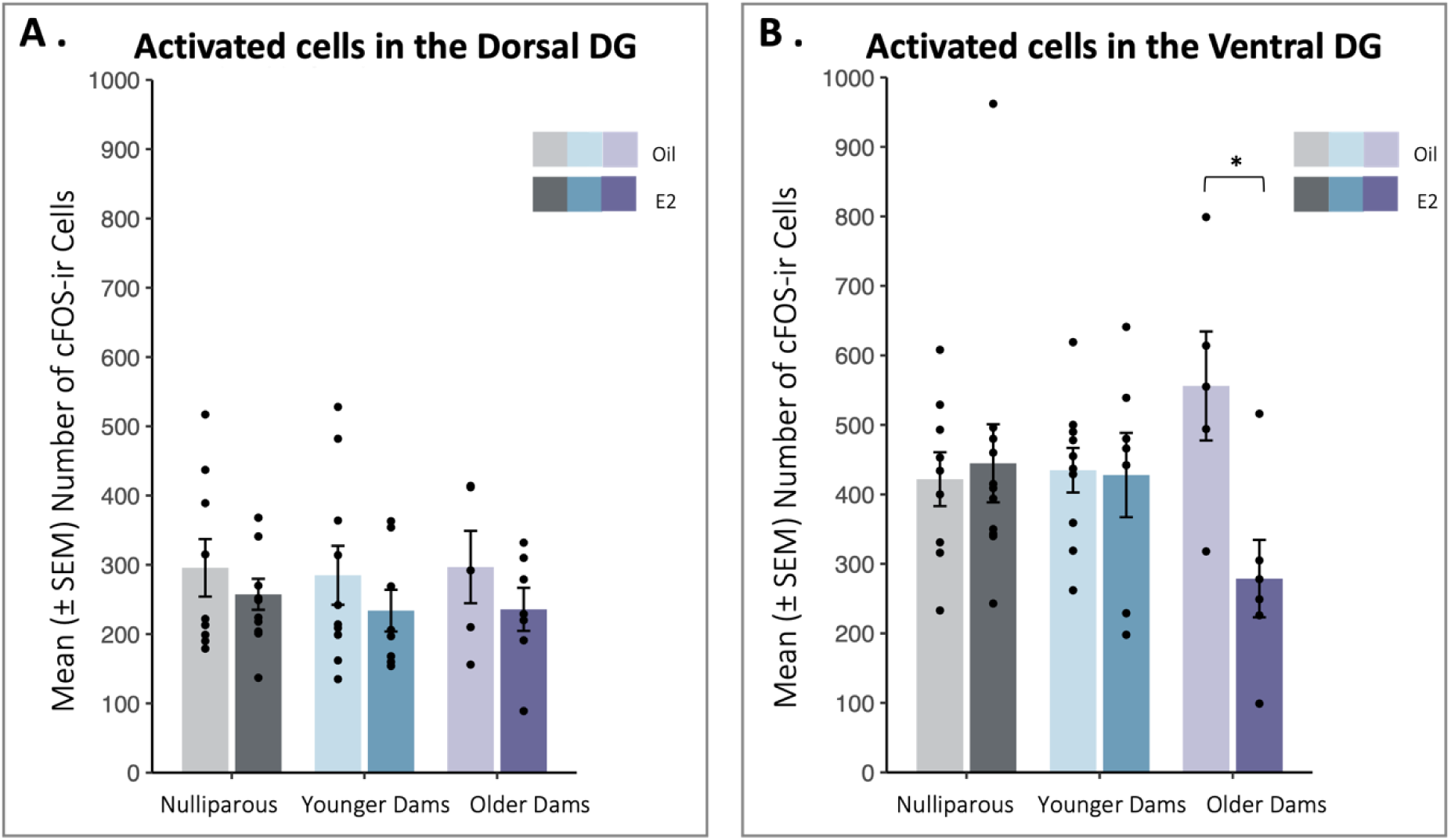
(A-B) Mean ± SEM (standard error of the mean) of number of c-FOS-ir cells in the (A) dorsal and (B) ventral DG. (A) In the dorsal DG, estradiol-treated animals show lower activation compared to oil-treated controls. (B) In the ventral dentate gyrus, estradiol lowers activation only in older dams. * Indicates p < 0.05. Abbreviations: DG = dentate gyrus; E2 = estradiol.

### 3.7 Estradiol lowers IL-4, IL-10, and IL-13 concentration based on the age of first pregnancy and lowers TNF-α concentration in the dorsal hippocampus regardless of parity

Estradiol lowered IL-13 concentrations in the dorsal hippocampus of younger dams (p=0.008, Cohen’s *d* = 1.298), but not in nulliparous (p=0.97) or older dams (p=0.42); [treatment by pregnancy interaction: F (2,47) = 3.25, p = 0.047, partial η^2^ = 0.121, Figure 5A]. There was also a main effect of treatment [main effect of treatment: F (1,47) = 9.37, p = 0.003, partial η^2^ = 0.166]. There were no significant main effects or interaction in the ventral hippocampus (all p’s > 0.109).

**Figure 5.**
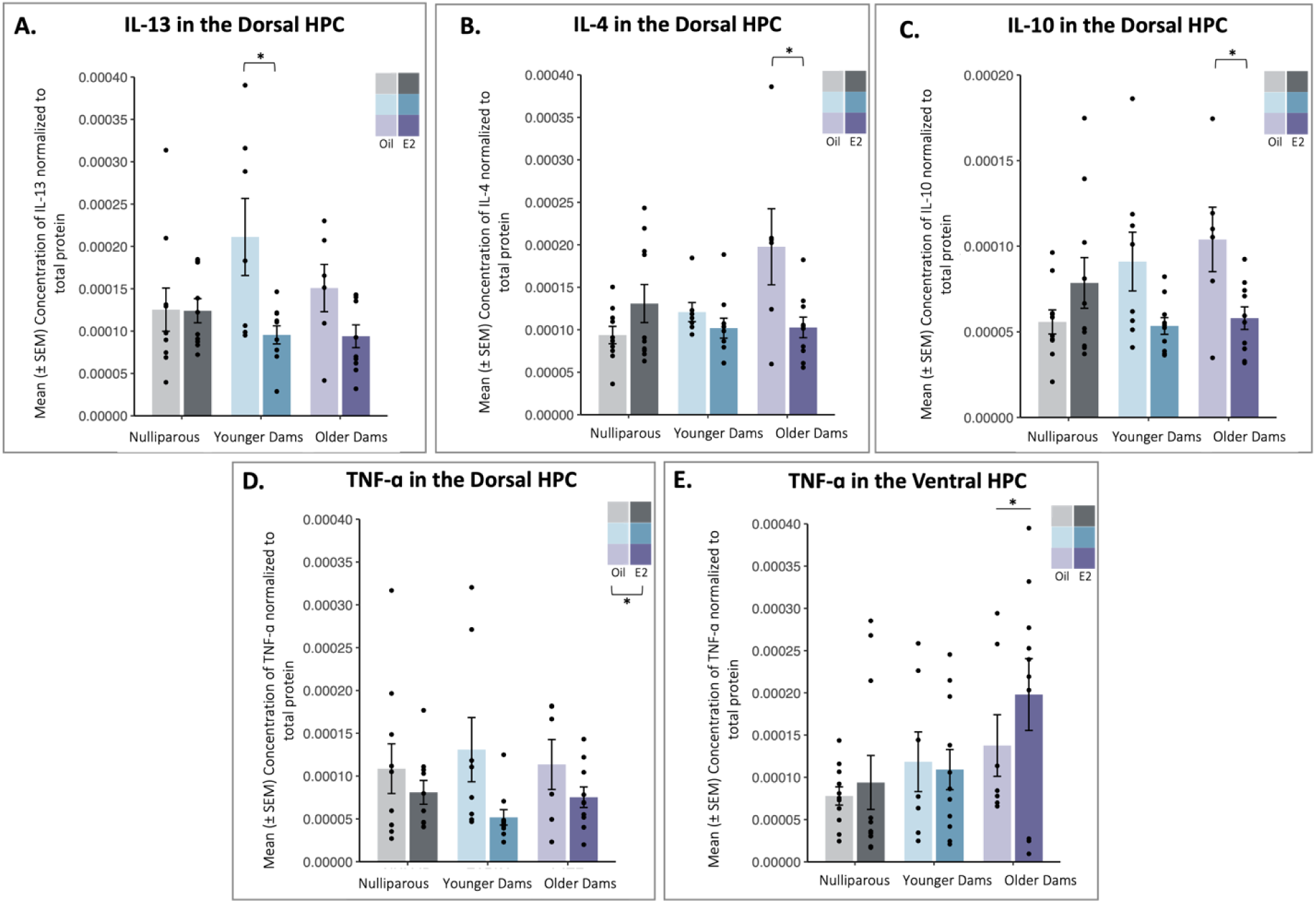
(A-E) Mean ± SEM (standard error of the mean) concentration of cytokines. (A) Concentration of IL-13 in the dorsal hippocampus (HPC). Estradiol lowered IL-13 concentration in younger dams. (B) Concentration of IL-4 in the dorsal HPC. Estradiol lowered IL-4 concentration in older dams (C) Concentration of IL-10 in the dorsal HPC. Estradiol lowered IL-10 concentration in older dams (D) Concentration of TNF-α in the dorsal HPC. Estradiol lowered TNF-α concentration regardless of parity. (E) Concentration of TNF-α in the ventral HPC. Older dams had a higher concentration of TNF-α in the ventral HPC. * Indicates p < 0.05. Abbreviations: HPC = hippocampus; E2 = estradiol.

Estradiol lowered the concentration of cytokine IL-4 in the dorsal hippocampus of older dams (p = 0.005, Cohen’s *d* = 1.154), but not in nulliparous (p = 0.650) or younger dams (p = 0.763) [treatment by pregnancy interaction: F (2,47) = 6.03, p = 0.005, partial η^2^ = 0.204, Figure 5B]. There was also a main effect of treatment [main effect of treatment: F (1,48) = 4.04, p = 0.048, partial η^2^ = 0.078, Figure 5B]. There were no significant main effects or interactions in the ventral hippocampus (all p’s > 0.445).

Estradiol also lowered the concentration of IL-10 in the dorsal hippocampus of older dams (p = 0.038, Cohen’s *d* = 1.286), but not nulliparous (p = 0.363) or younger dams (p = 0.173) [treatment by pregnancy interaction: F (2,48) = 5.42, p = 0.007, partial η^2^ = 0.184, Figure 5C]. There was also a main effect of treatment [main effect of treatment: F (1,48) = 4.49, p = 0.039, partial η^2^ = 0.085, Figure 5C]. There were no significant main effects or interactions in the ventral hippocampus (all p’s > 0.533).

In the dorsal hippocampus, estradiol treated rats had lower TNF-α concentrations than oil-treated controls, regardless of parity [main effect of treatment: F (1,48) = 6.706, p = 0.013, partial η^2^ = 0.123; Figure 5D]. In the ventral hippocampus, dams pregnant at an older age had higher concentration of cytokine TNF-α regardless of estradiol treatment [main effect of treatment F (1,52) = 4.605, p = 0.014, partial η^2^ = 0.150, Figure 5E]. There were no other main effects or interactions in the dorsal or ventral hippocampus (all p’s > 0.481 and 0.748 respectively).

## 4. Discussion

Here we examined the influence of estradiol and age of first pregnancy on various aging biomarkers in the hippocampus in middle aged rats. Estradiol, regardless of age of first pregnancy, reduced activation of new neurons, and reduced TNF-α concentrations in the dorsal hippocampus compared to vehicle-treated rats. An older age of first pregnancy reduced spatial memory, improved reversal learning, increased neurogenesis in the ventral hippocampus, and increased both activation and TNF-α concentration in the ventral DG compared to the other group regardless of estradiol treatment. For inflammatory markers, the age of first pregnancy also influenced the effects of estradiol. In the dorsal hippocampus, estradiol reduced IL-13 concentration in younger dams, but reduced IL-4 and IL-10 concentrations in older dams compared to other groups. In this study, we report that pregnancy history and/or estradiol exposure affected spatial memory and reversal learning in the Morris water maze, neurogenesis in the hippocampus, and inflammatory signaling in the hippocampus.

### 4.1 The age of first pregnancy influenced spatial memory and reversal learning

Surprisingly, we did not find any significant effect of reproductive history or chronic estradiol on acquisition in the water maze. Other studies have found that previous parity enhances initial acquisition of spatial working and reference memory in middle age (Barha et al., 2015; Galea et al., 2018; Talboom et al., 2008; Zimberknopf et al., 2011). It is not clear why our study found no effect of parity on Morris water maze learning; however, we did see that rats pregnant at an older age did show reduced distance to reach the hidden platform during reversal learning. This improved performance relative to the other groups was likely driven by the fact that these older dams did not perform above-chance during the spatial memory trial, unlike nulliparous and younger dams. As the reversal task is dependent on the prefrontal cortex, the increased neurogenesis in the ventral dentate gyrus of older dams may be related to their performance, as the ventral hippocampus projects to the frontal cortex (Dalton et al., 2016; Phillips et al., 2019). Previous work shows that increased TNF-α levels in the hippocampus is associated with impaired spatial-dependent memory (Takahashi et al., 2021). Our study found that older dams had increased levels of TNF-α in the ventral hippocampus compared to both younger dams and nulliparous animals. Thus, it is possible that increased TNF-α concentration in the older dams could be underlying the impaired performance of older dams on the probe trial.

### 4.2 An older age of first pregnancy increases hippocampal neurogenesis in middle age compared to a younger age of first pregnancy

In the present study we find that dams pregnant at an older age showed greater neurogenesis than dams pregnant at a younger age in the ventral dentate gyrus. In humans, a younger age of first pregnancy is associated with a trend in smaller brain volume (Hoekzema et al., 2017). Some previous studies have report that primiparity increases neurogenesis in middle age, but we find no significant difference between younger dams and nulliparous rats (Eid et al., 2019; Galea et al., 2018; Lee et al., 2023). However, this may be explained by the fact that all three of these studies, rats were pregnant at an older age than the younger dams in our experiment (At 7 months, 5 months, and 4 months respectively). Therefore, it may be the age of first pregnancy is contributing, at least in part, to these effects. As neurogenesis declines with age (Kuhn et al., 1996), the fact that previous parity is associated with higher levels of neurogenesis indicates that reproductive experience is likely neuroprotective. Our results, in part, align with ‘BrainAGE’ data in humans showing that females that have previously given birth have ‘younger looking’ brains compared to those who have not, but this study did not investigate the age of first pregnancy, and it could be a potential contributor to these results (de Lange et al., 2020).

We did not find significant effects of estradiol on neurogenesis, regardless of parity. This is somewhat inconsistent with previous work which finds that that 22 days of E2 reduced the number of DCX-expressing cells in multiparous animals but not nulliparous animals, and Premarin, an estrone-based MHT, reduced the number of immature neurons in the ventral dentate gyrus of middle-aged primiparous rats (Barha et al., 2015; Galea et al., 2018). The differences in our data could be explained by the number of pregnancies affecting the response to estrogens, different lengths of treatment, and different formulations of MHTs as well. This highlights the need for more investigations into how course, dose, and reproductive experience affect the brain’s response to estrogens.

### 4.3 Estradiol reduces activation of new neurons in the DG, regardless of age of first pregnancy and reduces ventral DG activation only in older dams

We show that rats treated with estradiol exhibited decreased cFOS expression in immature neurons in the dorsal dentate gyrus compared to vehicle-treated controls. We also found, contrary to previous work in younger females, that estradiol reduced activation in the dentate gyrus in the rats that got pregnant at an older age. Previous research has indicated that high levels of estradiol are associated with heightened activation of new-born neurons, immediate-early gene expression, and functional connectivity (Hidalgo-Lopez et al., 2020; McClure et al., 2013; Yagi et al., 2017, 2023). However, all of these studies were performed in young rodents or humans, compared to our findings in middle-aged rats. Furthermore, there is evidence that aging can alter responsiveness of neuronal activation to estradiol (Meyza et al., 2007). Therefore, estradiol could be increasing cFOS responsiveness or activation in younger but increasing it in older individuals, and further research is required to elucidate any potential mechanisms.

### 4.4 Estradiol reduces cytokine concentration in the dorsal hippocampus

In the dorsal hippocampus, estradiol affected the concentration of three cytokines depending on the age of first pregnancy. Estradiol decreased levels of IL-13 in younger dams, and decreased levels of IL-4 and IL-10 in older dams. These three cytokines are ‘anti-inflammatory’ and have very closely linked functions, and are secreted by Th2 type T-cells (KleinJan et al., 1999; Yssel & Groux, 2000). Although there are some previous studies that have investigated the effects of parity and inflammation and found no effect of parity on IL-13, IL-10, or TNF-α, they did not consider any hormonal treatment or age of first pregnancy, and also looked at circulating cytokines, whereas our study investigated cytokine levels in brain tissue (Duarte-Guterman et al., 2023; Eid et al., 2019).

Estradiol also decreased TNF-α expression in the dorsal hippocampus regardless of the age of first pregnancy. This is consistent with a previous body of work that indicates estradiol lowers the expression of proinflammatory cytokines (Azcoitia et al., 2019; Vegeto et al., 2006; Z. Zhang et al., 2018), including TNF-α in the hippocampus (W. Zhang et al., 2020). Our work aligns with previous studies indicating that decreased levels of estrogens in middle age are associated with higher levels of inflammation (Au et al., 2016; Salem, 2004).

In the ventral hippocampus, older dams showed increased concentration of TNF-α regardless of estradiol treatment. There is evidence that TNF-α imbalance can contribute to pregnancy complications such as pre-eclampsia, and human studies show older mothers (> 40 years) are at increased risk for these conditions (Kirwan et al., 2002; Sheen et al., 2020). It is possible that TNF-α is mediating risks to other processes as well, in a manner related to the age of first pregnancy, and furthermore, contributed to the impaired performance of older dams during the probe trial in our study.

## 5. Conclusion

In this study, we find that rats that were pregnant at an older age had more neurogenesis in the ventral hippocampus than those pregnant at a younger age rats. An older age of first pregnancy also affected memory and cognitive flexibility, and increased neurogenesis, activation of cells, and TNF-α concentration in the ventral dentate gyrus during middle age. We suggest that an older age of first pregnancy might be playing a beneficial role in brain aging by increasing neurogenesis and improving cognitive flexibility in middle aged rats. Estradiol alone, irrespective of the age of first pregnancy, decreased activation of new neurons and reduced TFN-α concentrations in the dorsal hippocampus when compared to oil-treated rats. Interestingly, estradiol’s effects also changed with the age of first pregnancy in some instances. Estradiol reduced the concentration of IL-13 in younger dams, but reduced concentrations of IL-4 and IL-10 in older dams only. Our results demonstrate not only that previous pregnancy, and the age of first pregnancy, can affect multiple brain parameters, but also that the effects of estradiol in middle age can vary with previous reproductive experience. This work converges with others to indicate that parity, and associated factors such as the number of pregnancies affect the trajectory of brain aging and health (Duarte-Guterman et al., 2019, 2023; Galea et al., 2018; Puri et al., 2023). With an aging global population, our work suggests that parity, and the age of first pregnancy should be taken into consideration when investigating the effects of estrogens in middle and older age. This area of research remains significantly understudied, and there is an urgent need to further examine the influences of previous reproductive experience and exogenous hormones on brain aging.

## Conflict of Interest

The authors have no conflict to declare.

## Acknowledgements

We gratefully thank Tallinn Splinter, Rebecca Rechlin, Dr. Travis Hodges, and Muna Ibrahim for their help with animal care procedures, as well as Anne Cheng and Alice Chan for going above and beyond in their work as animal care technicians at the Center for Disease Modeling at the University of British Columbia. We also would like to thank Lydia Huntsman and Harleen Hans for helping collect and quantify behavioral data.

## Funding Sources

This work was funded by CIHR Project Grant 148662 to LAMG, and University of British Columbia Four Year Fellowship to TAP.

## SUPPLEMENT

**Figure A.1.**
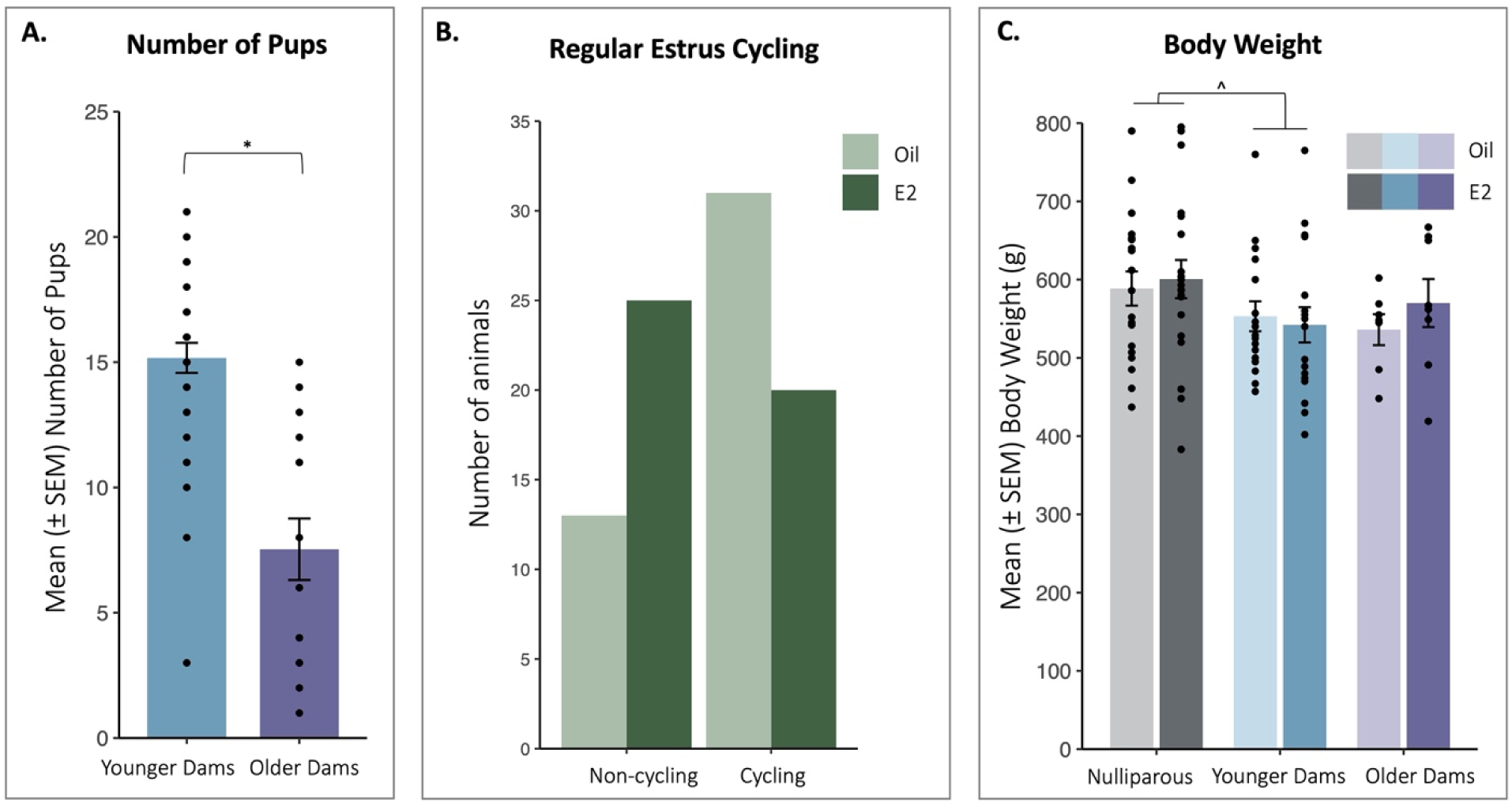
(A) Mean ± SEM (standard error of the mean) of the number of pups each dam had before litters were culled. Younger dams had more pups than older dams. (A) Frequency of the number of dams that were either cycling or not based on estradiol treatment. Estradiol treated animals are more likely to have an irregular estrus cycle than those treated with oil vehicle. (B) Mean ± SEM body weight in rats by age of first pregnancy. Trend towards nulliparous rats being heavier than younger dams approached significance. * Indicates p < 0.05. ^ indicates 0.05 < p < 0.10.

**Table A.1.** Mean ± SEM time (in seconds) per 10-minute observation block spent engaging in active maternal behaviors (such as nursing and licking of pups) and off nest behaviors during postnatal days 2-8.

| Group | Nursing/Licking | Off-Nest |
| --- | --- | --- |
| Younger Dams | 87.59 ± 3.91 | 42.51 ± 4.30 |
| Older Dams | 88.16 ± 4.62 | 31.39 ± 4.63 |

## Notes

### Competing Interest Statement

The authors have declared no competing interest.

